# Evaluating the Efficacy of Propofol Against Isoniazid-Induced Seizures: A Comparative Study with Diazepam and Pyridoxine in Swiss Albino Mice

**DOI:** 10.1101/2025.09.17.676905

**Authors:** Prinay Sohal, Kanchan Gupta, Sandeep Kaushal, Michael E. Mullins

**Affiliations:** Department of Pharmacology, Dayanand Medical College & Hospital, Ludhiana, Punjab (141001), India; Division of Medical Toxicology, Department of Emergency Medicine, Washington University School of Medicine, MSC 8072-17-06, 660 S. Euclid Avenue, Saint Louis, Missouri 63110-1010, USA

**Author notes:** Corresponding author: Prinay Sohal, MBBS, Dayanand Medical College & Hospital, Ludhiana, India. **Funding disclosure Statement** This research did not receive any funding from funding agencies in the public, commercial, or not-for-profit sectors. **Authors’ contributions (CRediT statement)** Conceptualization: Michael E. Mullins, Prinay Sohal; Methodology: Prinay Sohal, Kanchan Gupta, Sandeep Kaushal; Investigation: Prinay Sohal, Kanchan Gupta, Sandeep Kaushal; Formal analysis: Prinay Sohal, Michael E. Mullins; Writing – original draft: Prinay Sohal; Writing – review & editing: Michael E. Mullins, Prinay Sohal, Kanchan Gupta, Sandeep Kaushal; Supervision: Michael E. Mullins.

**Keywords:** Isoniazid toxicity, seizures, propofol, pyridoxine, diazepam

## Abstract

**Objective:** Acute isoniazid toxicity causes refractory seizures due to pyridoxine deficiency-mediated gamma-aminobutyric acid depletion. While pyridoxine is the established antidote, its limited availability in emergency settings necessitates alternative treatments. This study aimed to evaluate the anticonvulsant efficacy of propofol, a GABA A receptor agonist and N-methyl-D-aspartate receptor antagonist, compared to pyridoxine and diazepam in isoniazid-induced seizures in mice.

**Methods:** Thirty-two Swiss albino mice were pretreated with: (1) normal saline (10 mL/kg), (2) pyridoxine (300 mg/kg), (3) propofol (50 mg/kg), or (4) diazepam (2.5 mg/kg), followed by isoniazid (300 mg/kg) after 30 minutes. All drugs were administered intraperitoneally. Seizure latency, duration, time to death, and survival were monitored for 120 minutes. Outcomes were analyzed using Kaplan-Meier/log-rank tests and ANOVA with Tukey’s post-hoc test.

**Results:** Isoniazid induced seizures and 100% mortality in the control group. Pyridoxine did not prevent seizures or improve survival (8/8 seizures and deaths). Propofol failed to improve seizure frequency, latency, duration, mortality, or time to death (8/8 seizures and deaths). Diazepam significantly reduced seizure frequency and mortality to 1 of 8 mice (p < 0.001), and delayed seizure onset.

**Conclusion:** Pretreatment with either propofol or pyridoxine failed to protect against isoniazid-induced seizures and death in mice. In contrast, diazepam was effective in preventing both seizures and death. Further studies are needed to investigate different propofol administration protocols and alternative animal models.

## INTRODUCTION

Isoniazid (isonicotinic acid hydrazine, commonly known as INH) is a bactericidal drug used in the treatment and prophylaxis of tuberculosis [1–3]. Isoniazid overdose causes seizures that are often refractory to standard anticonvulsants [4–7]. Isoniazid is one of the leading causes of toxicant-induced seizures and accounted for 6% of the cases in a report from the California Poison Control System [8]. Without prompt and effective treatment, Isoniazid toxicity can progress to status epilepticus, metabolic acidosis, and coma, and may be fatal [9].

Isoniazid inhibits the pyridoxine phosphokinase which converts pyridoxine (vitamin B_6_) into its active form pyridoxal-5-phosphate [1,2]. Pyridoxal 5’-phosphate is a necessary cofactor for conversion of glutamate to gamma-aminobutyric acid (GABA) [10–12]. The imbalance of high concentrations of glutamate (an excitatory neurotransmitter) and low concentration of GABA (an inhibitory neurotransmitter) leads to seizures. Benzodiazepines, the first-line anticonvulsants for seizures of toxic origin, require the presence of GABA, and therefore, fail to produce a meaningful response in a GABA-deficient state [2,4]. Currently, high-dose intravenous pyridoxine supplementation is the only effective treatment available in cases of acute isoniazid toxicity [2]. Pyridoxine corrects this imbalance to normalize GABA concentrations. Pyridoxine also restores the effectiveness of benzodiazepines. However, the large dose required (up to 5 g) exceeds supplies in many hospital pharmacies [13–17].

Propofol is a sedative-hypnotic agent with anesthetic properties widely used in emergency departments, operating theaters and intensive care units [18,19]. Propofol uniquely opens GABA-A receptors coupled chloride channels, even in the absence of GABA [20,21]. Propofol is also an N-methyl-D-aspartate (NMDA) receptor antagonist, which reduces excitatory transmission in the brain during an ongoing seizure [22]. Its wide availability, and unique actions make it a compelling potential treatment for isoniazid induced seizures [20–24]. We sought to compare propofol to diazepam and pyridoxine in an experimental animal model of severe isoniazid poisoning.

## METHODS

A total of 32 Swiss albino mice, weighing 20-25 g each, were used in this study. Each individual mouse was considered an experimental unit. All animals were sourced from the Central Animal House at Dayanand Medical College & Hospital, Ludhiana, India. The mice were housed in groups of five in standard polycarbonate cages with husk bedding, and free access to a standard rodent chow diet and water. The animal room was maintained at a temperature of 22 ± 2°C with a 12-hour light/dark cycle.

All experimental procedures were approved by the Institutional Animal Ethics Committee (IAEC) and conducted in accordance with the Committee for Control and Supervision of Experiments on Animals (CCSEA) guidelines for the care and use of laboratory animals (425/Po/ReBi/S/01/CPCSEA).

We obtained isoniazid (gift sample, PGI, Chandigarh, India), diazepam (LORI, Neon Laboratories Ltd., India), propofol (MCT-ROF, Neon Laboratories Ltd., India), and pyridoxine (pyridoxine hydrochloride, Benadon, Bayer Pharmaceuticals Ltd., India) for use in this study. We dissolved isoniazid and pyridoxine in normal saline immediately before use. The doses were selected based on previous studies [25–29].

We assessed the effect of propofol on isoniazid neurotoxicity using a previously published study design [25]. We divided the animals into four treatment groups of eight each and pretreated with different reagents. Group 1 received normal saline 10 mL/kg by intraperitoneal (IP) injection. Group 2 received pyridoxine 300 mg/kg IP. Group 3 received propofol 50 mg/kg IP. Group 4 received diazepam 2.5 mg/kg IP.

All animals then received isoniazid IP at the neurotoxic dose (300 mg/kg) to induce neurotoxicity 30 minutes after the initial drug administration [30]. We observed the mice for 120 minutes post-isoniazid injection for seizure activity. We recorded the latency to onset and duration of convulsions for each mouse. We recorded the number of mice surviving at the end of the observation period and the latency to death of those that died.

We used Statistical Kingdom (Melbourne, Australia; http://www.statsking.com) to analyze survival data using the log-rank test and to generate Kaplan–Meier survival curves for time to seizure and time to death. Seizure duration data were analyzed using JASP software (Version 0.19.3, JASP Team, Amsterdam, The Netherlands) and are expressed as mean ± standard error of the mean (mean ± SEM). Statistical analysis was conducted using one-way ANOVA followed by Tukey’s Honestly Significant Difference (HSD) post hoc test. A p-value of < 0.05 was considered statistically significant.

## RESULTS

Intraperitoneal injection of isoniazid produced convulsions and death in all 8 mice in the control group, with all animals experiencing seizures and dying within 60 minutes (Fig. 1). Compared to control, animals in the pyridoxine and propofol groups had similar median latency times to seizure, mean seizure durations, and median latency times to death (Table 1, Fig. 1). All animals in the propofol group eventually died, indicating no survival benefit. In contrast, diazepam was best in both latency to seizure and latency to death. It prevented seizure in 7 of 8 mice and delayed seizure onset in the remaining mouse. Only 1 of 8 of animals had convulsions after diazepam, and 7 of 8 had seizure-free survival to 120 min. The animal that seized and died had a seizure latency of 108 min and death latency of 109.2 min (both approximately double the median latencies in all other groups).

**Table 1:** Effects of propofol, diazepam, and pyridoxine on isoniazid-induced seizure activity and mortality in mice.

| Treatment Group | Seizure (%) | Seizure Latency (min)<br>medians (range) | Seizure Duration (s)<br>mean ± SEM | Death (%) | Latency to Death (min)<br>medians (range) |
| --- | --- | --- | --- | --- | --- |
| Control | 8/8 (100%) | 57.5 min (47-65 min) | 21.5 ± 2.1 | 8/8 (100%) | 60.5 min (48-70 min) |
| Propofol | 8/8 (100%) | 51.5 min (41-72 min) | 18.3 ± 3.1 | 8/8 (100%) | 52.5 min (44-80 min) |
| Diazepam | 1/8 (12.5%)** | 120 min (108-120 min)* # | 20<sup>##</sup> | 1/8 (12.5%)** | 120 min (109-120 min)* # |
| Pyridoxine | 8/8 (100%) | 41.5 min (31-54 min) | 24.2 ± 8.0 | 8/8 (100%) | 43.5 min (36-84 min) |

\* $p < 0.001$ by log rank test.
\*\* $p < 0.001$ compared to control by Fisher's exact test for frequencies for diazepam.
#Diazepam data censored at 120 min.

**Fig. 1.**
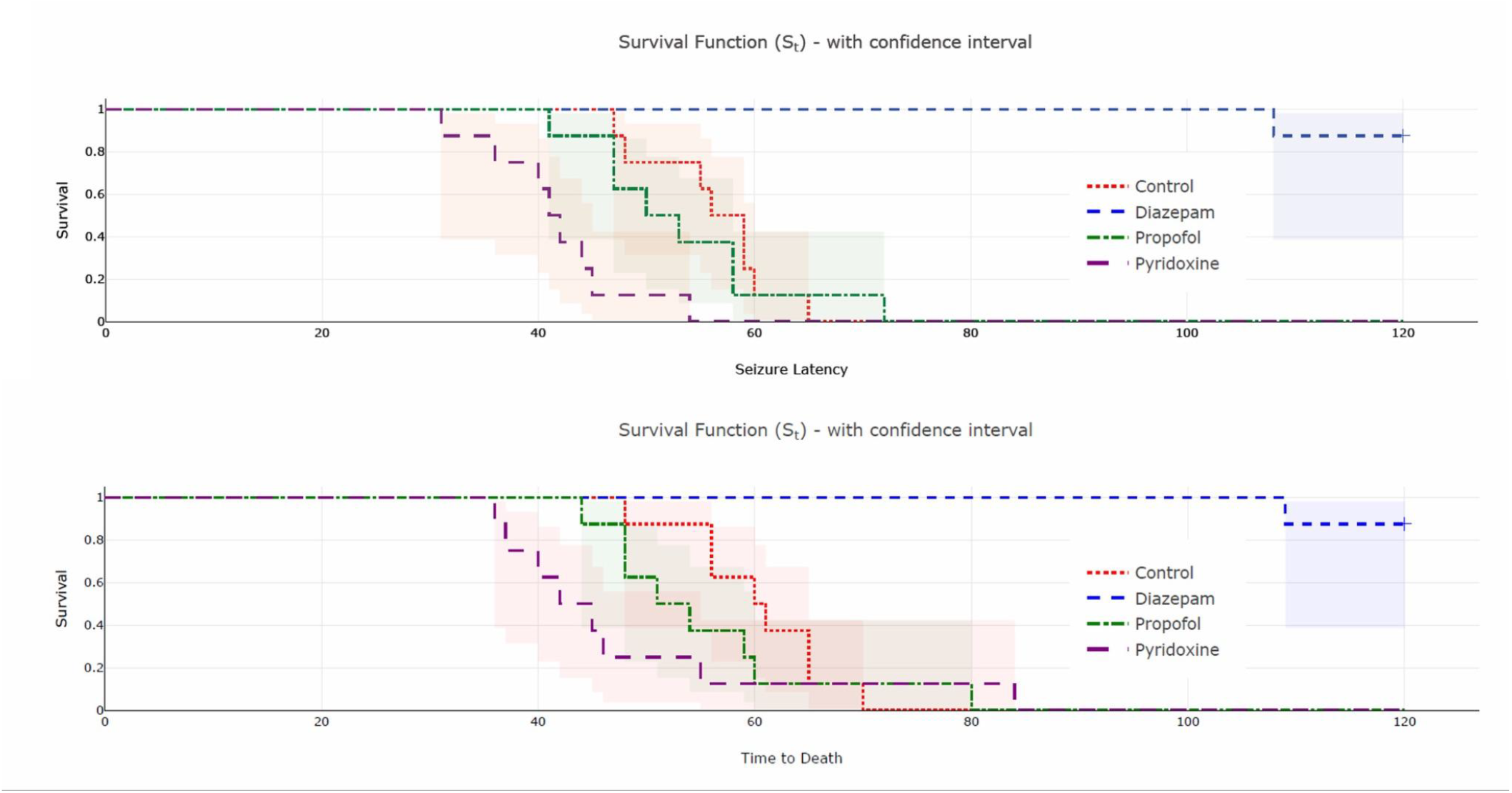
Kaplan-Meier survival curves showing seizure latency (top) and time to death (bottom) following isoniazid-induced seizures in mice treated with different agents. Shaded areas indicate 95% confidence intervals

## DISCUSSION

Pyridoxine is the established antidote for isoniazid-induced seizures, as it replenishes GABA by restoring pyridoxal 5’-phosphate (PLP) synthesis [2,6,31]. However, in our study pyridoxine (300 mg/kg, IP) not only failed to prevent seizures or death but paradoxically shortened seizure latency. This is a finding contrary to human clinical experience, as pyridoxine both treats and prevents isoniazid-induced seizures in humans [2]. The finding is, however, congruent with previous studies showing ineffectiveness of pyridoxine in isoniazid-induced seizures in mice [25,32,33]. Studies also suggest that PLP depletion may not be the primary mechanism for isoniazid neurotoxicity in mice and indicate other mechanisms for isoniazid seizures that remain unaffected by pyridoxine repletion [32,33].

We hypothesized that propofol, a direct GABA-A agonist and NMDA antagonist [20–22], might bypass isoniazid-induced GABA deficiency. It did not delay seizure onset or improve survival, indicating that propofol was not able to counteract isoniazid’s neuronal injury in our murine model. Propofol has a very short duration of action, its elimination half-life in the brain being less than 10 minutes in mice [34,35]. The experimental procedure involved administering only a single dose of propofol 30 minutes prior to isoniazid and then observing for 120 minutes. The drug likely metabolized and cleared from brain neural tissue well before the completion of the observation period, rendering any assessment of its sustained anticonvulsant properties fundamentally unreliable. A better protocol would involve maintaining therapeutic brain concentrations throughout the entire observation period. The anticonvulsant action of propofol in rats is temporary, after which rebound seizures tend to occur [36].

Unlike in this pre-treatment model, propofol use in clinical settings usually occurs after the seizure onset [37,38]. Propofol also has a proconvulsant action, which may have contributed to the results [39]. One of the reasons for hypothesized superiority of propofol in comparison to diazepam was the PLP depletion mechanism of isoniazid neurotoxicity in human beings which, as mentioned earlier, is not the case in mice [32,33]. Therefore, no further comment can be made on its efficacy in isoniazid neurotoxicity without further experiments in other species models, with underlying mechanisms closer to human pathogenesis.

Diazepam was the most effective agent in our model, which contrasts with its limited efficacy in treating isoniazid toxicity in humans. The depletion of PLP resulting in GABA depletion is the primary mechanism of isoniazid neurotoxicity in human beings and explains the failure of benzodiazepines in isoniazid-poisoned patients. As this mechanism is not at play in mice [32,33], diazepam can use the GABA present in neuronal clefts to exert its anticonvulsant effect in this model. Diazepam also inhibits excitotoxicity independently of GABA which could also explain its superiority to other GABA modulatory anticonvulsants [40–42]. Previous studies showed that the effectiveness of anticonvulsant medications depends more on their capacity to prolong the latency of seizures than on their ability to prevent convulsions [25] and diazepam’s long duration of action allows maintaining adequate concentration in brain during the 120 min observation period [43].

Our study has several additional limitations. First, the use of albino mice may not fully reflect human isoniazid toxicity. Mice may have mechanisms of isoniazid neurotoxicity other than GABA deficiency to explain the discrepancies in the effects of pyridoxine and diazepam. Second, we evaluated each drug using only one dose concentration and one administration route. This is particularly problematic for propofol which has a short duration of action. The single-dose approach may not reflect how propofol would be delivered in actual clinical practice. Continuous or repeated dosing, and drug administration by intravenous route might have yielded different results.

We also relied on behavioral seizure scoring without electroencephalographic monitoring, possibly missing subclinical seizures and underestimating drug effects. Biochemical analysis of brain tissue for oxidative stress and antioxidant concentrations could also be employed to give deeper insight into the neuroprotective effects of the tested drugs against seizure-induced brain damage.

Future studies should use animal models more similar to humans, such as rats, and explore IV doses and infusions of propofol to maintain adequate concentration during the observation period. Incorporating electroencephalography and biochemical analysis would provide a more comprehensive assessment of drug effect.

In conclusion, propofol did not significantly delay seizure onset or improve survival. In contrast, diazepam delayed seizures and prevented mortality. Pyridoxine, the standard antidote, was also ineffective in delaying seizures and preventing mortality.

## Acknowledgements

The authors thank the technical staff of the Central Animal House and the Department of Pharmacology, Dayanand Medical College & Hospital, for their kind assistance. We are also grateful to Dr. Samir Malhotra (PGI, Chandigarh, India) for providing the isoniazid used to induce toxicity, and to Dr. Vir Vikram (CT University, Ludhiana, India) for his guidance on animal handling.

